# Genome-wide DNA methylation changes in human spermatogenesis

**DOI:** 10.1101/2023.10.27.564382

**Authors:** Lara M. Siebert-Kuss, Verena Dietrich, Sara Di Persio, Jahnavi Bhaskaran, Martin Stehling, Jann-Frederik Cremers, Sarah Sandmann, Julian Varghese, Sabine Kliesch, Stefan Schlatt, Juan M. Vaquerizas, Nina Neuhaus, Sandra Laurentino

## Abstract

Sperm production and function require the correct establishment of DNA methylation patterns in the germline. Here, we examined the genome-wide DNA methylation changes during human spermatogenesis and its alterations in disturbed spermatogenesis. We found that spermatogenesis is associated with remodeling of the methylome, comprising a global-decline in DNA methylation in primary spermatocytes followed by selective remethylation, resulting in a spermatid-specific methylome. Hypomethylated regions in spermatids were enriched in specific transcription factor binding sites for DMRT and SOX family members and spermatid-specific genes. Intriguingly, while SINEs displayed differential methylation throughout spermatogenesis, LINEs appeared to be protected from changes in DNA methylation. In disturbed spermatogenesis, germ cells exhibited considerable DNA methylation changes, which were significantly enriched at transposable elements and genes involved in spermatogenesis. We detected hypomethylation in SVA and L1HS in disturbed spermatogenesis, suggesting an association between the abnormal programming of these regions and failure of germ cells progressing beyond meiosis.

## Introduction

There is growing evidence that the establishment of the male germ cell methylome is not restricted to embryonic development, but continues in adulthood during spermatogenesis (Oakes *et al*, 2007; Langenstroth-Röwer *et al*, 2017; Di Persio *et al*, 2021a; El Omri-Charai *et al*, 2023; Huang *et al*, 2023). Especially in the early phases of meiosis, when replication and recombination occur, the genome is hypomethylated. This event is highly conserved between mouse and human and has been hypothesized to result from a delay in DNA methylation maintenance (Gaysinskaya *et al*, 2018; Huang *et al*, 2023). Still, there is limited information on DNA methylation during human spermatogenesis or whether disturbances in this process impact spermatogenesis and sperm function.

Epigenetic remodeling, which includes reprogramming of the methylome, is essential for cell-fate decisions in mammals (Klemm *et al*, 2019; Markenscoff-Papadimitriou *et al*, 2020; Izzo *et al*, 2020). Recently, it has been demonstrated that the impaired function of enzymes involved in the DNA methylation machinery results in a range of disturbances to spermatogenesis, including complete sterility (Bourc’his *et al*, 2001; Zamudio *et al*, 2015; Barau *et al*, 2016; Karahan *et al*, 2021; Dura *et al*, 2022). In most cases of severe male infertility, sperm output is drastically reduced and the etiology is unknown (Tüttelmann *et al*, 2018). In particular, cryptozoospermia is characterized by a drastic decline in germ cells going through meiosis and an accumulation of the most undifferentiated spermatogonia (Di Persio *et al*, 2021b). Previously, we reported aberrant DNA methylation in bulk germ cells of men displaying cryptozoospermia (Di Persio *et al*, 2021a), which led us to hypothesize that aberrant DNA methylation could be the underlying cause. However, the extent to which the different germ cell types carry altered DNA methylation, the genomic regions affected, and their involvement in post-meiotic germ cell decline, remain to be assessed.

Especially during meiosis, the genome heavily depends on the suppression of transposable elements (TEs). This is usually achieved by epigenetic mechanisms such as gene-silencing histone modifications, piRNA machinery, and DNA methylation (Zamudio & Bourc’his, 2010, Zamudio et al 2015). Thus, the suppression of TEs, such as long and short-interspersed nuclear elements (LINE/SINE), is crucial to maintain genome integrity in the germline (Schaefer *et al*, 2007; Vaissiere *et al*, 2008; Zamudio & Bourc’his, 2010; Dong *et al*, 2019), which, when lost, leads to sterility in mice (Barau *et al*, 2016; Vasiliauskaitė *et al*, 2018). Differential methylation at TEs or dysfunctional TE silencing pathways have been linked to male infertility (Bourc’his & Bestor, 2004; Carmell *et al*, 2007; Aravin *et al*, 2007; Heyn *et al*, 2012; Urdinguio *et al*, 2015). Importantly, different TEs have different modes of DNA methylation acquisition (Fukuda et al 2022), and evolutionary younger (SINE-VNTR-Alus (SVAs)) TEs are protected from genome-wide methylome erasure during development (Seisenberger *et al*, 2012; Kobayashi *et al*, 2012; Gkountela *et al*, 2015; Guo *et al*, 2015), indicating that these elements are especially hazardous to the genome and must be protected via DNA methylation. The protective role of DNA methylation at different TEs in germ cells during spermatogenesis and its association with male infertility, in particular during the hypomethylated phase of meiosis, remains to be elucidated.

By performing whole methylome analysis on pure human male germ cell fractions, we uncovered that human spermatogenesis is associated with epigenetic remodeling of the methylome. We found that the global decline in DNA methylation in primary spermatocytes is followed by selective remethylation in specific regions in spermatids, suggesting that the hypomethylated primary spermatocyte genome is not exclusive a transient side effect of DNA replication but is required for the establishment of a spermatid-specific methylome. We found significant differences in DNA methylation in germ cells of infertile men, particularly occurring in intergenic regions and TEs, and demonstrate that different TEs are differentially reprogrammed during spermatogenesis. Our study increases current evidence on the role of DNA methylation changes during human spermatogenesis, and points to an association between altered DNA methylation and spermatogenic failure.

## Results

### Primary spermatocytes exhibit genome-wide reduced DNA methylation levels

To investigate genome-wide DNA methylation changes during human spermatogenesis, we performed whole-genome enzymatic methyl-sequencing (EM-seq) on isolated human male germ cells. We obtained undifferentiated spermatogonia (2C/MAGEA4^+^/UTF1^+^/DMRT1^−^; Undiff), differentiating spermatogonia (2C/MAGEA4^+^/UTF1^−^/DMRT1^+^; Diff), primary spermatocytes (double-diploid cells; 4C), and spermatids (haploid cells; 1C) from men with normal spermatogenesis (controls, CTR) (Fig. 1A, Supplementary Fig. S1A). We captured an average of 28,049,115 CpG sites per sample, including 26,861,322 CpGs with a minimum coverage of 5 reads, which were the ones considered for further analyses.

**Fig. 1:**
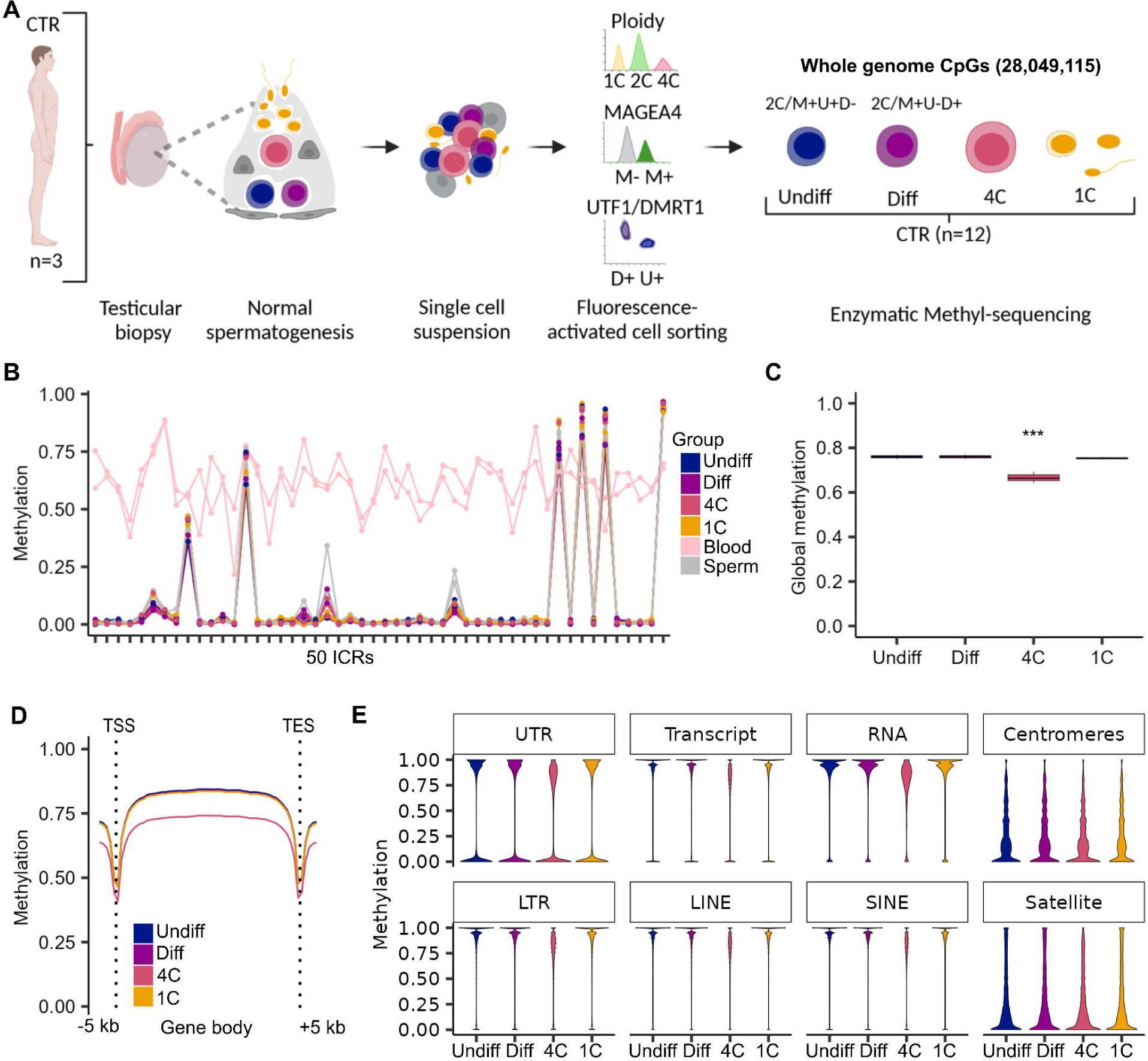
Primary spermatocytes exhibit genome-wide reduced DNA methylation levels. **A** Schematic illustration on the retrieval of whole genome methylome data from germ cells of samples with normal spermatogenesis (control, CTR). CpGs refer to the mean CpG number captured by enzymatic methyl-sequencing in all germ cell fractions (n=12). **B** Lineplot depicts the methylation in 50 imprinted control regions (ICRs) for each CTR sample and germ cell type compared to published blood and sperm samples (Laurentino *et al*, 2020). **C** Box plots display the mean global DNA methylation levels. Statistical tests: ANOVA test followed by Tukey-HSD-test: *** < 0.001 of 4C compared to all other germ cells. Data are represented as median (center line), upper/ lower quartiles (box limits), 1.5 x interquartile range (whiskers). **D** Lineplot shows mean methylation levels per group across gene bodies divided into 50 intervals (bins) and 5 kb upstream and downstream of the transcriptional start sites (TSSs) and transcriptional end sites (TESs). **E** Violin plots represent methylated CpGs across different genomic compartments. Panel A was created with BioRender.com. See also Supplementary Fig. S1.

To guarantee that all germ cell fractions were free of somatic DNA, we compared the methylation of published sperm-soma differentially methylated regions (DMRs) (Leitão *et al*, 2020) in our germ cell fractions with those published for sperm and blood samples (Laurentino *et al*, 2020). We confirmed male germ cell-specific methylation in the isolated cell fractions (Supplementary Fig. S1B). Interestingly, the methylation patterns of 50 known maternally and paternally methylated imprinted control regions (ICRs) (Monk *et al*, 2018) were also similar to those found in sperm and had the same methylation pattern in all germ cell types (Fig. 1B, Supplementary Table S1). Analysis of global DNA methylation levels across spermatogenesis revealed comparably high average levels (>74%) of DNA methylation in undifferentiated spermatogonia, differentiating spermatogonia, and spermatids (Fig. 1C), which is consistent with the characteristically high DNA methylation levels in male germ cells (Greenberg & Bourc’his, 2019). Intriguingly, the global DNA methylation levels in primary spermatocytes were significantly lower, with a mean of 66% (Fig. 1C), compared to all other germ cell types. To elucidate whether the DNA hypomethylation in primary spermatocytes occurs randomly in the genome or is specific to particular genomic features, we analyzed the DNA methylation in gene bodies and 5000 bp up- and downstream of the transcriptional start (TSS) and end sites (TES). Our data showed that the decreased DNA methylation in primary spermatocytes, which was evident at TSS and TE, and was particularly prominent in gene bodies (Fig. 1D). Deeper analysis of the different genomic compartments revealed a decline in DNA methylation in primary spermatocytes that occurred across untranslated regions (UTRs), transcripts, and RNA repeats. DNA methylation also declined at long terminal repeats (LTRs), including the TEs LINEs and SINEs, indicating a genome-wide demethylation in primary spermatocytes (Fig. 1E). In contrast, there was no change in DNA methylation at centromeric and satellite regions, which overall display low DNA methylation levels in all germ cell types evaluated.

### Differentially methylated regions during spermatogenesis are enriched at SINE repeats

In order to identify regions that change their DNA methylation during spermatogenesis, we compared the methylomes of the different germ cell-types (coverage ≥ 5, p-value ≤ 0.05 (metilene), difference ≥ 20 %) using metilene and camel (Wöste *et al*, 2020). Applying stringent filtering, we identified a total of 16,000 DMRs throughout the different germ cell types (Fig. 2A, Supplementary Table S2). We found the fewest DMRs between undifferentiated and differentiating spermatogonia (64 DMRs, mean Δ methylation =24%), indicating a high level of similarity between the methylomes of the two spermatogonial subpopulations. The most DMRs were detected between spermatogonia (undifferentiated and differentiating) and primary spermatocytes (5212 and 5487 DMRs, mean Δ methylation = 22% and 23%). When we compared the cell types corresponding to the least and most differentiated cell types, namely undifferentiated/differentiating spermatogonia and spermatids, we found the most extreme changes in DNA methylation (1516 and 1001 DMRs, mean Δ methylation = 41% and 36%). Analyses on the intersection of DMRs with genes or promoters revealed that 53 - 70% are associated with genes and 3 - 9% with a promoter (Fig. 2B). To investigate the features of the DMRs we analyzed their CpG enrichment and length. In comparison to other cell type comparisons that had an average of 8 CpGs and a length of 340 bp, we found that DMRs obtained from the comparisons that included primary spermatocytes were the longest (average 975 bp) and the most enriched in CpGs (average 14 CpGs) (Fig. 2C+D). We asked whether certain chromosomes showed an enrichment in DMRs. To this end, we normalized the number of DMRs per chromosome size (bp) (Piovesan *et al*, 2019) and scaled for the number of DMRs per group comparison. We found that DMRs were similarly distributed across the different chromosomes in all comparisons, except for the 64 undifferentiated vs differentiating spermatogonia DMRs, which showed a peak on the X chromosome (Fig. 2E). Moreover, we sought to elucidate whether the DMRs are significantly enriched for certain genomic features. Intriguingly, enrichment analysis showed that particularly DMRs of the undifferentiated spermatogonia vs spermatids comparison, were significantly enriched (corrected p-value ≤ 0.00019, z-score >0) at CpG shelves, 3’UTRs, and exons, indicating changes in DNA methylation in these regions might be characteristic of the process of spermatogenesis (Fig. 2F). We found reduced global DNA methylation levels of LINEs and SINEs exclusively in primary spermatocytes (Fig. 1E). For all DMRs between the different germ cell types, we found a significant enrichment only at SINEs (Fig. 2F), suggesting a role for DNA methylation changes in SINEs during spermatogenesis. In contrast, we found an underrepresentation compared to what would be expected by chance (z-score < 0) of promoters, CpG shores and islands in DMRs of the differentiating vs primary spermatocyte comparison, putatively indicating the methylation levels of these regions are maintained during the entry into meiosis. Interestingly, LINE elements were underrepresented in the linear DMR comparisons (i.e. Undiff vs Diff, Diff vs 4C, 4C vs 1C) as well as between the undifferentiated spermatogonia vs primary spermatocyte comparison, indicating a preferential DNA methylation retention at LINE repeat regions during spermatogenesis.

**Fig. 2:**
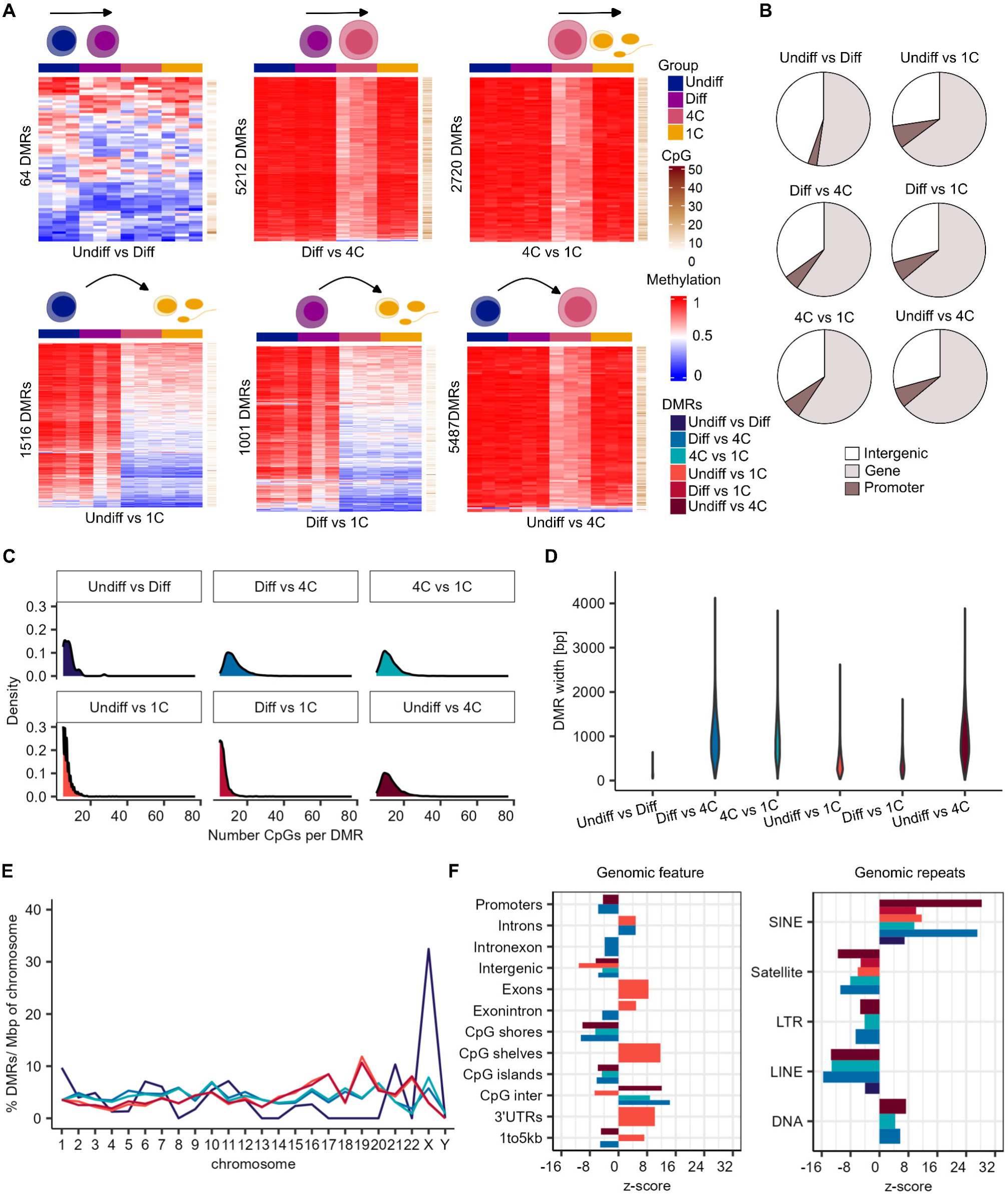
Differentially methylated regions during spermatogenesis are enriched at SINE repeats. **A** Heatmaps display methylation values and CpG numbers of the differentially methylated regions (DMRs) of all germ cell type comparisons. **B** DMRs are associated with genes and promoters. **C** Frequency of CpG number per DMR within the different group comparisons. **D** Violin plot depicts distribution of the DMR width in each group comparison. **E** Distribution of DMRs per chromosome scaled for chromosomal size (basepairs) and normalized by their total count within one group. **F** Enrichment of DMRs for general genomic features and genomic repeats. Positive and negative enrichments are indicated by z-score. Displayed annotations, p < 0.00019 by permutation tests. Color coding of the group comparisons are depicted in panel A.

To unravel the putative regulatory role of DNA methylation changes in biological processes in the respective germ cell types, we investigated the nearby genes with a regulatory region within the identified DMRs. We found that the DMRs of all comparisons overlapped a putative regulatory region of 29 to 1685 genes (Supplementary Table S3). Within these genes, gene ontology (GO) analysis revealed an enrichment of distinct GO terms in biological processes (BP) and molecular functions (MF), such as regulation of cation channel activity (Undiff vs Diff), and GTPase regulatory activity (Undiff vs 1C) (Supplementary Fig. S2) indicating changes in DNA methylation during spermatogenesis associates with specific cellular functions.

### Hypomethylated regions in spermatids are enriched for TF binding sites

Local methylation changes can identify potential regulatory regions in developing germ cells (Kubo *et al*, 2015). To address whether regulatory transcription factor (TF) binding sites are subject of changes in DNA methylation during spermatogenesis, we analyzed the DMRs for their enrichment in TF binding motifs by applying the Hypergeometric Optimization of Motif Enrichment (HOMER) analysis. This analysis revealed that particularly the DMRs of the spermatogonia (undifferentiated and differentiating) vs spermatids comparisons were enriched for motifs recognized by TFs such as *DMRT1*, *DMRT6/ DMRTB1,* and *SOX6* (Fig. 3A). To assess whether these TFs are expressed in the respective germ cell types, in which we found differential methylation of their motifs, we analyzed their expression during spermatogenesis in our published scRNA-seq dataset (Di Persio *et al*, 2021b). We found germ cell type-specific expression of *DMRT1*, *DMRTB1,* and *SOX6* in differentiating spermatogonia, spermatocytes, and spermatids, respectively (Fig. 3B), suggesting the change in DNA methylation of the identified motifs is a feature of spermatogenesis.

**Fig. 3:**
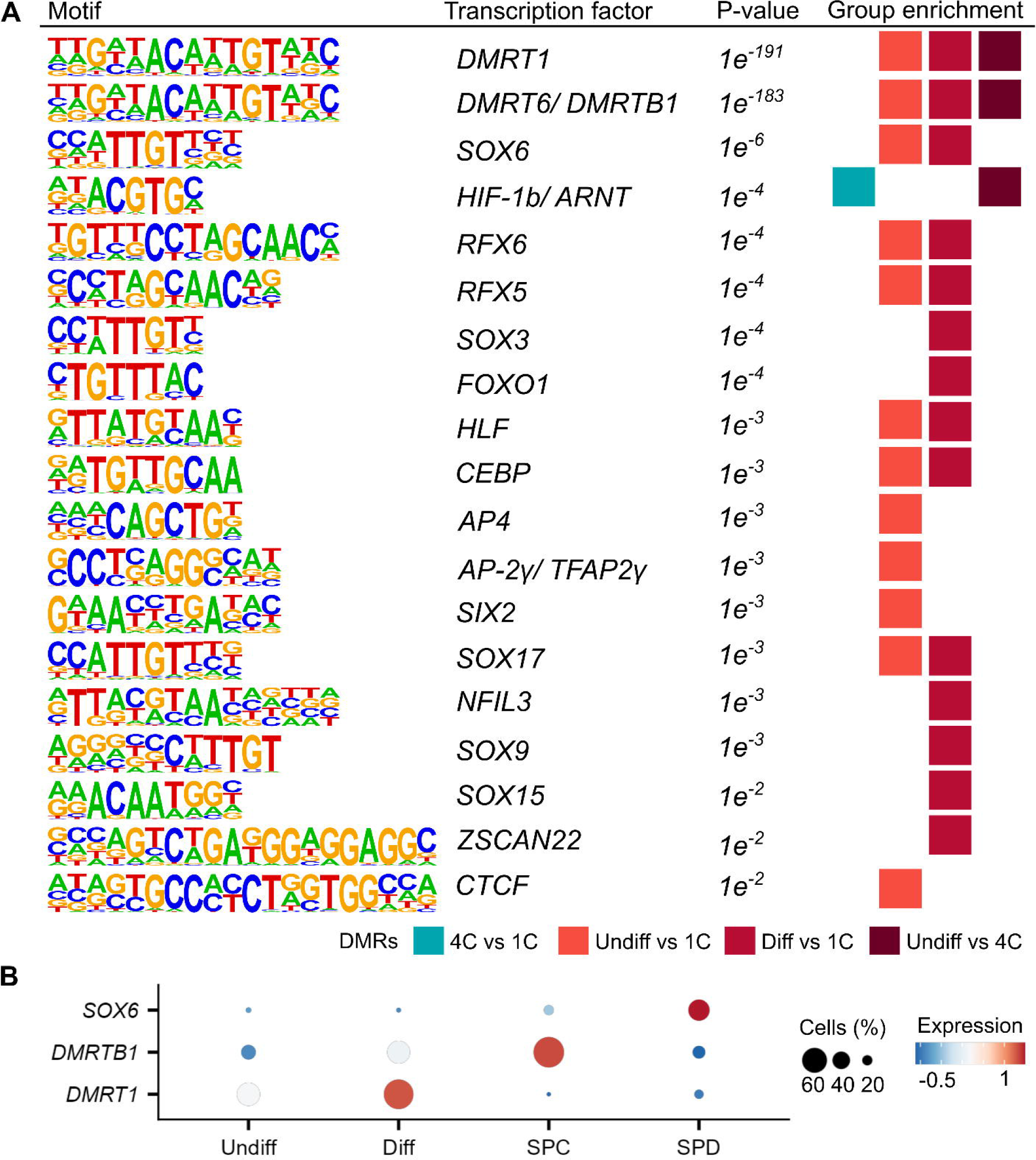
Hypomethylated regions in spermatids are enriched for specific TF binding sites. **A** Depicted are the enriched sequences of known motifs identified by HOMER. HOMER analysis was run for all DMR comparisons and significant results are displayed (p-value < 0.01, FDR <0.05). **B** Dot plots show the average single cell expression (Di Persio *et al*, 2021b) of the top 3 transcription factors (TF) with enriched motifs among the DMRs identified with HOMER. SPC = spermatocytes, SPD = spermatids.

### Germ cell type-specific expression of the DMR associated genes

Low or unmethylated regions frequently mark accessible chromatin regions (Stadler *et al*, 2011). As we found that the methylomes of spermatids, when compared to undifferentiated spermatogonia, have the highest number of hypomethylated DMRs and are enriched for TF binding sites, we hypothesized that the hypomethylated DMRs mark functionally important regions for spermatids. To this end we evaluated whether the DMR associated genes (overlapping a gene body) of the undifferentiated spermatogonia vs spermatids comparison have specific expression in spermatids. Indeed, we identified 24 highly spermatid-specific genes (e.g. *RAD21L1, KLK11, FEM1B, CLPB, NRDC*) among the hypomethylated DMRs (Fig. 4A). Intriguingly, and in line with the association of gene expression and gene body methylation (Ball *et al*, 2009), we found 41 spermatogonial-specific genes (e.g. *SERPINE2, BLVRA, TJP1, YPEL2, TCF3*) that have a hypermethylated DMR in their gene bodies in undifferentiated spermatogonia (Fig. 4A). When we analyzed the DMR associated genes of other group comparisons, we observed that the majority of germ cell type-specific genes are hypermethylated (Supplementary Fig. S3A) and only a minority of them are hypomethylated in the respective germ cell types (Supplementary Fig. S3B).

**Fig. 4:**
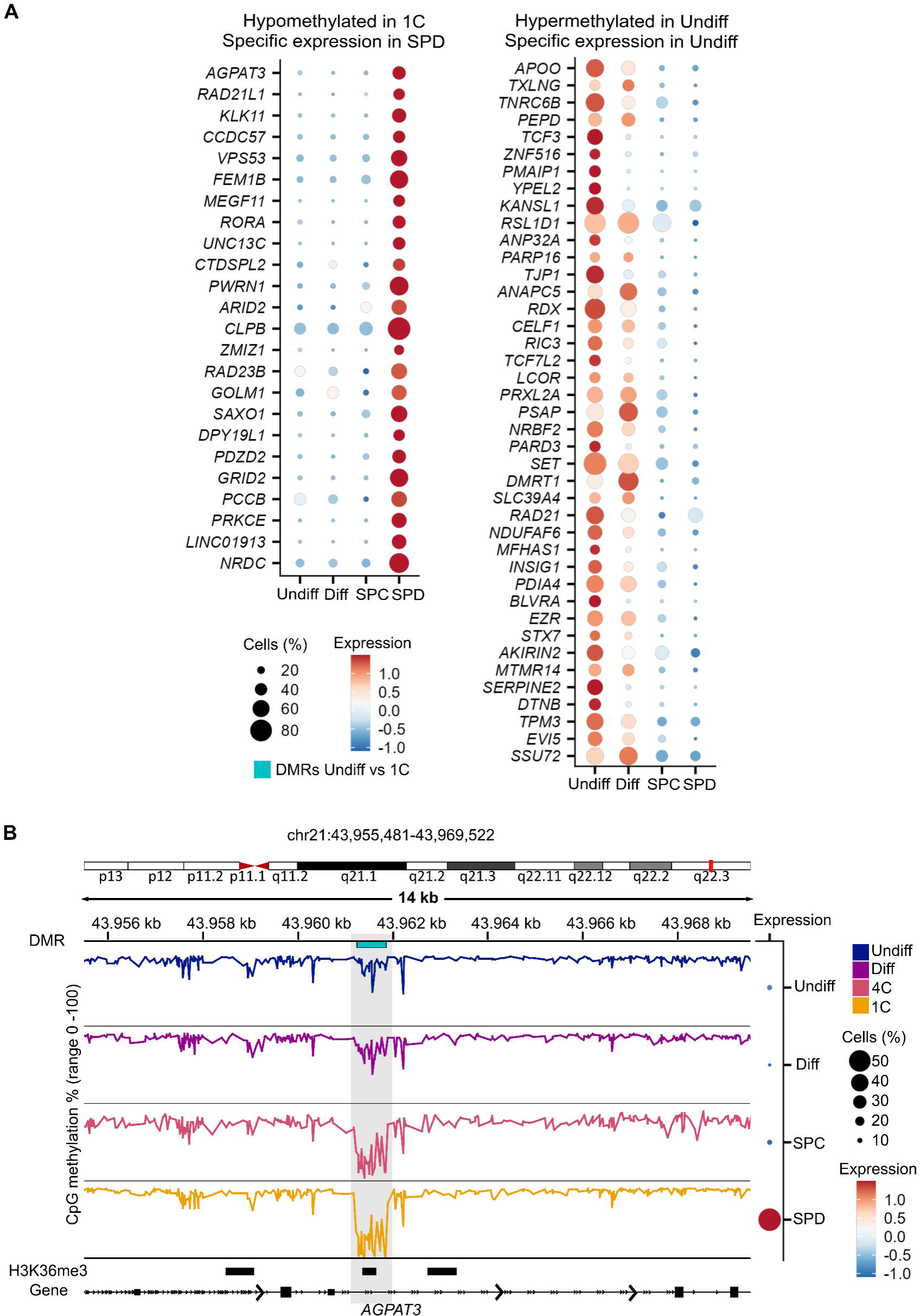
Hypomethylated regions in the spermatid methylome mark spermatid-specific genes. **A** Single cell expression (Di Persio *et al*, 2021b) of hypomethylated DMR associated genes with specific expression in spermatids and hypermethylated DMR associated genes with specific expression in undifferentiated spermatogonia of the undifferentiated spermatogonia vs 1C DMRs. **B** Example of the DMR methylation within the *AGPAT3* locus that is hypermethylated in Undiff and Diff and hypomethylated in 4C and 1C and specifically expressed in SPD. H3K36me3 histone modification data (GSE40195) in human sperm is also shown. SPC = spermatocytes, SPD = spermatids. See also Supplementary Fig. S3.

Retained nucleosomes in human sperm were shown to be enriched at gene regulatory regions (Hammoud *et al*, 2009; Brykczynska *et al*, 2010). To examine whether the hypomethylated DMRs of the undifferentiated spermatogonia vs spermatids comparison mark regulatory active regions in sperm, we analyzed the overlap of the hypomethylated DMRs with retained histone marks in sperm. Indeed, overlap analysis with publicly available datasets on human sperm histones (GSE40195) revealed that ∼17% of DMRs overlapped with a retained histone mark in sperm (Supplementary Table S4). The active histone mark H3K36me3 overlaid 10% of the hypomethylated DMRs in spermatids. Although H3K36me3 is associated with gene body methylation to maintain gene expression stability (Sharda & Humphrey, 2022), we found H3K36me3 marked a hypomethylated region in *AGPAT3*, which is specifically expressed in spermatids (Fig. 4B).

### Disturbed spermatogenesis displays methylome changes at TEs and spermatogenesis genes

Aberrant DNA methylation was found in sperm of infertile men, particularly in imprinted genes (Marques *et al*, 2004; Poplinski *et al*, 2010; Kläver *et al*, 2013; Kuhtz *et al*, 2014; Laurentino *et al*, 2015; Urdinguio *et al*, 2015). To confidently associate changes in DNA methylation with male infertility, it is important to exclude effects of differential methylation, especially in imprinted genes, due to potential contamination with somatic DNA. Accordingly, it remains to be elucidated whether genome-wide changes in germ cells of infertile patients occur. To this end, we analyzed methylomes of germ cells isolated from testicular tissues of men diagnosed with cryptozoospermia (CZ) (Fig. 5A). Ploidy analysis confirmed the characteristic cryptozoospermic-phenotype (Di Persio *et al*, 2021b), consisting of a decreased proportion of spermatids in comparison to the CTR samples (Fig. 5B). Based on cell numbers (Supplementary Table S5), we were able to generate methylomes of undifferentiated spermatogonia, differentiating spermatogonia, and primary spermatocytes from these samples. Quality control (Leitão *et al*, 2020; Di Persio *et al*, 2021a) indicated the presence of somatic DNA in CZ-1 (Supplementary Fig. 4A+B), therefore we exclude all fractions from this patient from further analyses. The other two samples were free of somatic DNA and, except for one secondary ICR (MKRN3:TSS-DMR) that showed an isolated increase in DNA methylation (average of 35%), showed no overall imprinting errors (Supplementary Fig. S4C, Supplementary Table S6).

**Fig. 5:**
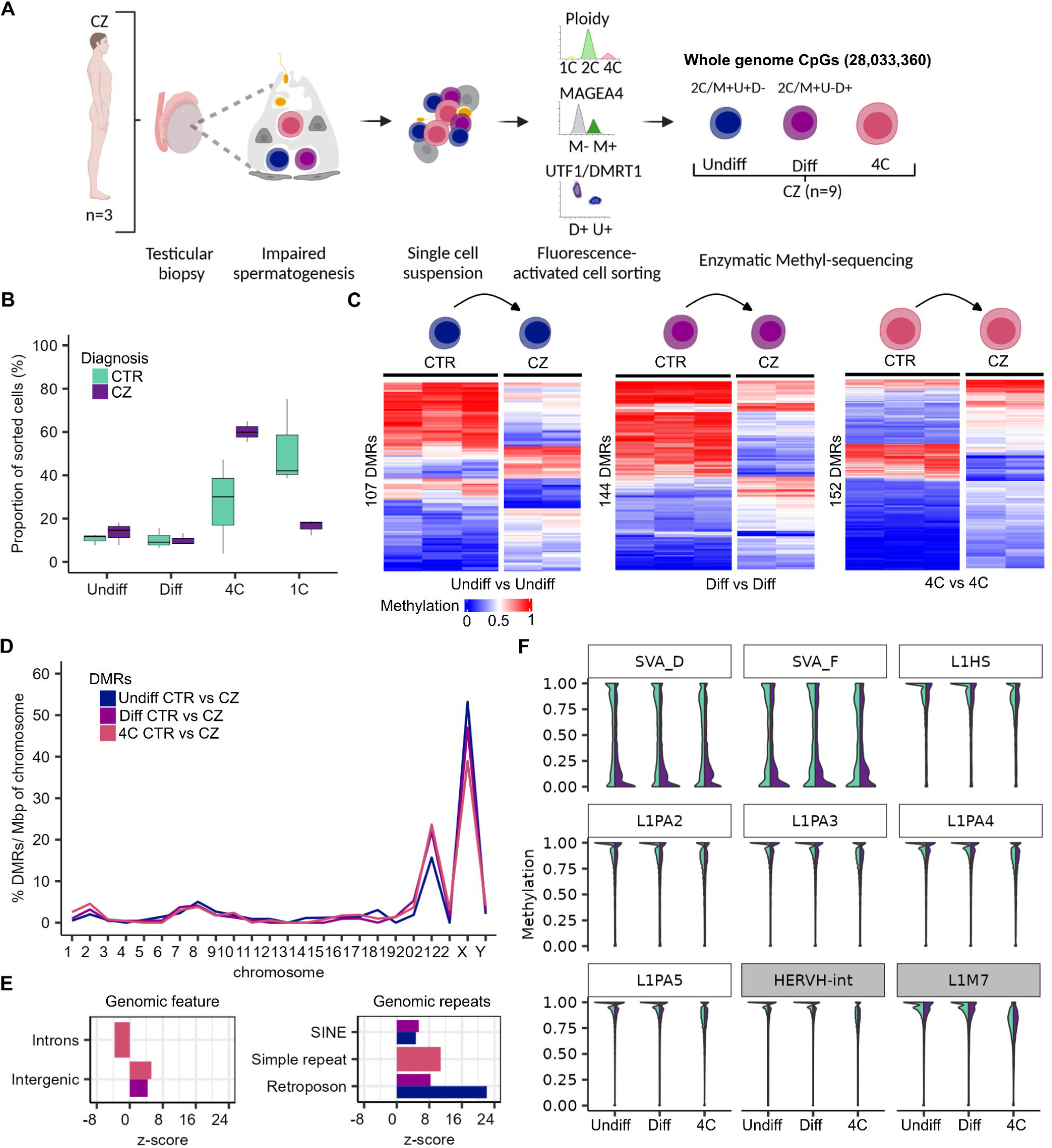
Disturbed spermatogenesis displays methylome changes at TEs and spermatogenesis genes. **A** Schematic illustration on the retrieval of whole genome methylome data of germ cells from samples with disturbed spermatogenesis (cryptozoospermia, CZ). **B** Box plots show the proportion of cell types among the sorted cells in the CTR and CZ patients. Data are represented as median (center line), upper/ lower quartiles (box limits), 1.5 x interquartile range (whiskers). **C** Heatmaps display methylation values of the differentially methylated regions (DMRs) between CTR and CZ of the same cell type (Undiff vs Undiff, Diff vs Diff, 4C vs 4C). **D** Distribution of the CTR/CZ DMRs per chromosome scaled for chromosomal size (basepairs) and normalized by their total count within one group. **E** Enrichment of CTR/CZ DMRs for functional general genomic regions and genomic repeats. Positive and negative enrichments are indicated by z-score. Displayed annotations, p < 0.00019 by permutation tests. **F** Violin plots showing the CpG methylation of evolutionary younger (white boxes: L1Hs, L1PA2-5, and SVA_D/F) and older (grey boxes: HERVH-int and L1M7) TEs in CTR and CZ germ cells. Color coding of the group comparisons are depicted in panel B. Panel A created with BioRender.com. See also Supplementary Fig. S4 and S5.

To examine whether regions exhibit differential methylation in cryptozoospermic patients (n=2) compared to control patients (n=3), we analyzed potential DMRs between undifferentiated spermatogonia, differentiating spermatogonia, and primary spermatocytes. We applied the same DMR filtering as for the control germ cell DMRs (coverage ≥ 5, p-value ≤ 0.05 (metilene), difference ≥ 20 %) but, due to the smaller sample size, we used stricter filtering parameters for the methylation range within the cryptozoospermic group (range ≤ 15%) to filter out DMRs potentially influenced by the genetic background. We found 107, 144, and 152 DMRs (mean Δ methylation = 31-37%) in undifferentiated spermatogonia, differentiating spermatogonia, and primary spermatocytes between CTR and CZ samples (Fig. 5C). We found few overlapping DMRs between all comparisons (Supplementary Table S7). The DMRs were overall enriched in chromosomes 21 and X (Fig. 5D) and significantly enriched in intergenic regions and at retroposons (Fig. 5E), indicating that altered DNA methylation patterns in cryptozoospermic germ cells are predominantly affecting these regions. Particularly hypomethylation of evolutionary young TEs have been associated with decreased fertility in mice (Karahan *et al*, 2021). Therefore, we examined whether we find changes in DNA methylation in evolutionary younger (SVA D/F, L1Hs, and L1PA2-5) and older TEs (HERVH-int and L1M7) between control and cryptozoospermic patients. We found that spermatogonia (undifferentiated and differentiating) and primary spermatocytes of cryptozoospermic men had a decrease in methylation in SVA D/F TEs (Fig. 5F). When we analyzed the methylation levels of different TEs in each control and cryptozoospermic sample, we found that the hypomethylation in TEs of the cryptozoospermic group is driven by sample CZ-2, which showed hypomethylation not only in SVA D/F, but also in SVA B/C as well as L1HS, L1PA2, HERVH-int (Supplementary Fig. S5). Interestingly, SVA A TEs were hypermethylated in both cryptozoospermic samples and in all control samples.

In bulk germ cells of cryptozoospermic samples, changes in DNA methylation were associated with functionally relevant genomic regions (Di Persio *et al*, 2021a). Our analysis revealed that CTR/CZ DMRs are associated with putative regulatory regions of 29, 39, and 38 genes in undifferentiated spermatogonia, differentiating spermatogonia, and primary spermatocytes, respectively (Supplementary Table S8), which were significantly enriched for genes involved in differentiation processes (e.g. cell morphogenesis involved in differentiation) (Supplementary Fig. S4D). As we found 29 - 40 % of DMRs overlap a gene body (Supplementary Fig. S4E), we asked whether they associate with germ cell type-specific gene expression, potentially pointing to a relevant function for spermatogenesis. Gene expression analysis showed that 25 - 33% DMR associated genes overlap with genes specifically expressed by spermatogonia (*AL157778.1, PCDH11X, EDA, ANKRD33B, AC087354.1, DACH2, VPS37D*), spermatocytes (*HSPBAP1, FOXK1, ARHGAP33, NBPF1, SLCO3A1*), or spermatids (*CCDC200, AL589935.1, ADARB2, SNHG27*) (Supplementary Fig. S4F). In summary, although we could not detect overall changes at ICRs, we found that cryptozoospermic germ cells exhibit changes in DNA methylation particularly occurring at evolutionarily younger TEs of the SAV familiy and at spermatogenic genes.

## Discussion

The establishment of the germ cell methylome extends beyond prenatal development (Oakes *et al*, 2007; Langenstroth-Röwer *et al*, 2017; Di Persio *et al*, 2021a; El Omri-Charai *et al*, 2023; Huang *et al*, 2023). In this study, we used whole methylome sequencing of germ cells to identify changes in DNA methylation occurring during human spermatogenesis. This included demethylation in primary spermatocytes followed by remethylation of specific regions, resulting in a unique spermatid-specific DNA methylation pattern. Furthermore, we identified changes in the DNA methylation of germ cells from infertile men particularly affecting intergenic regions and TEs.

Previous studies have reported that the male germline genome becomes hypomethylated during meiosis. They hypothesized that this decline in methylation is due to a delay in DNA methylation maintenance (Gaysinskaya *et al*, 2018) and is associated with meiotic recombination (Huang *et al*, 2023). Here, we show that the demethylation occurring in human male meiosis affects gene bodies and genomic repeats similarly, but not centromeres and satellite regions. Our data supports the hypothesis of a replication-dependent passive DNA demethylation in early meiosis resulting in a hypomethylated genome in spermatocytes (Gaysinskaya *et al*, 2018; Huang *et al*, 2023). This phase is followed by an increase in average DNA methylation to levels similar to those prior to meiosis. However, careful examination of regions showing differential methylation demonstrates that discrete regions remain protected from re-methylation in spermatids. This indicates that genome-wide re-methylation in spermatids is not an outcome of DNA maintenance machinery restoring CpG methylation after meiosis, but instead evidences the establishment of a highly specific DNA methylation pattern in spermatids.

Our findings lead us to hypothesize that a functional relationship exists between the establishment of a spermatid-specific methylome and gene expression during spermatogenesis. This notion was reinforced by our finding that hypomethylated regions in spermatids retain active histones in sperm. These regions correspond to genes specifically expressed by spermatids. Our hypothesis is further corroborated by analogous results in rodents, which demonstrate that the spermatid methylome undergoes differential methylation during spermiogenesis and is enriched for regulatory binding sites when compared to spermatogonial stem cells (Kubo *et al*, 2015; El Omri-Charai *et al*, 2023).

During normal human spermatogenesis, we showed that primary spermatocytes were the only germ cell type displaying a global decrease in DNA methylation at LTRs, LINEs, and SINEs. The silencing of TEs is crucial for maintaining genome integrity and usually achieved by epigenetic modifications, including repressive histone modifications and DNA methylation (Zamudio & Bourc’his, 2010). In line with this, our data showed that LINEs are significantly underrepresented among the regions with differential methylation, indicating that the methylation status of LINEs is protected in adult male germ cells. In contrast, we found that DNA methylation at SINEs significantly changes in all germ cell types during spermatogenesis, indicating that these regions are not under the same tight regulation as LINEs. Consistent with our data on genome-wide DNA methylation changes at SINE elements, a recent study found varying levels of SINE methylation in sperm, specifically at promoters of spermatogenic genes (Lambrot *et al*, 2021). Considering our data and the role of TEs in the regulation of genes (Zamudio & Bourc’his, 2010; Sasaki *et al*, 2008), we hypothesize that SINE methylation might play a regulatory role in human spermatogenesis.

Mice deficient for genes belonging to the DNA methylation machinery display male infertility or sterility (Bourc’his *et al*, 2001; Zamudio *et al*, 2015; Barau *et al*, 2016; Karahan *et al*, 2021; Dura *et al*, 2022). In humans, the hypothesis that male infertility is associated with aberrations in DNA methylation was first explored by numerous studies reporting epimutations in the sperm of infertile men (Åsenius *et al*, 2020), a notion that was somewhat challenged by our findings that somatic DNA artifacts and gene variants may have confounded previous studies (Leitão *et al*, 2020). However, more recently we identified consistent DNA methylation changes in testicular germ cells, but not in sperm, of infertile men, indicating that DNA methylation abnormalities during spermatogenesis might in fact be associated with disturbed spermatogenesis (Di Persio *et al*, 2021a). Here, we have expanded on this by identifying clear changes in DNA methylation at specific steps of spermatogenesis in germ cells of infertile men. Intriguingly, although these changes were germ cell type-specific, the affected regions were consistently enriched in TEs and predominantly localized on chromosomes 21 and X, chromosomes associated with monogenic disorders and male germ cell differentiation (Soumillon *et al*, 2013; Sangrithi *et al*, 2017; Ernst *et al*, 2019). In one of the cryptozoospermic samples, we specifically found hypomethylated SVA elements, together with hypomethylated L1HS, which is required to express SVA elements (Raiz *et al*, 2012). This finding shows similarities with a study demonstrating an association between hypomethylated young TEs and male infertility in mice (Karahan *et al*, 2021). DNA methylation is one of the main repressors of TE expression (Schaefer *et al*, 2007; Vaissiere *et al*, 2008; Zamudio & Bourc’his, 2010; Dong *et al*, 2019). Therefore, we hypothesize that the protection of evolutionary young TEs of the SVA family is disrupted in this patient’s germline leading to genomic instability due to the expression of SVAs and a phenotype of disturbed spermatogenesis. However, limited material prevents us from substantiating this at transcriptional level. Our findings on aberrant TE methylation in germ cells of men with cryptozoospermia are, nevertheless, supported by studies showing that loss of DNA methylation at TEs results in sterility in mice (Barau *et al*, 2016; Vasiliauskaitė *et al*, 2018). Failure in TE silencing affects germline gene expression in mice (Vasiliauskaitė *et al*, 2018) and, especially during meiosis, can lead to meiotic arrest (Bourc’his & Bestor, 2004; Carmell *et al*, 2007; De Fazio *et al*, 2011). In cryptozoospermic patients, some seminiferous tubules show complete spermatogenesis while others display spermatogenic arrest (Di Persio *et al*, 2021b). This heterogeneous phenotype could be caused by epimutations in the germline, in contrast to genetic variations that usually lead to complete phenotype penetrance. In line with this, we found that numerous infertility-related DMR associated genes are expressed during spermatogenesis (e.g. *VPS37D, FAM9B, SLCO3A1, CCDC200*).

In conclusion, our findings provide an unprecedented view of genome-wide DNA methylation changes during human spermatogenesis, highlighting the role of DNA methylation, particularly at TEs, during this process, and indicating a potential role for altered TE methylation in the etiology of human male infertility.

## Acknowledgments

We thank the Cologne Center for preparing the sequencing libraries and performing the sequencing. We appreciate the excellent technical support in patient material collection, histological evaluation of the testicular tissues and hormone measurement by Nicole Terwort, Sabine Forsthoff, Heidi Kersebom, Elke Kößer and Elisabeth Lahrmann as well as help from the andrology lab (CeRA). Panels Fig. 1A and Fig. 5A were created with BioRender.com.

This study was carried out within the frame of the Deutsche Forschungsgemeinschaft (DFG, German Research Foundation) sponsored Clinical Research Unit ‘Male Germ Cells’ (CRU326, project number LA 4065/3-2 & NE 2190/3-2 to NN & SL) & Innovative Medical Research (IMF) of the Medical Faculty of the University of Münster (to SDP).

## Author contributions

Study design: NN, SL; Conceptualization: LS, VD, NN, SL; Patient counseling: JC, SK; Data curation: LS, JB, MS, JC, SK; Formal analysis: LS, VD, SDP, MS; Investigation: LS, VD, SDP, JMV, NN, SL; Supervision: StS, SaS, JV, SK, JMV, NN, SL; Funding acquisition: SDP, JMV, NN, SL; Writing original draft: LS, VD, NN, SL; Writing review & editing: all authors.

All authors approved the final version of the manuscript.

## Materials & Methods

### Ethical approval

Patients included in this study underwent surgery for (microdissection) testicular sperm extraction (mTESE; n=6) at the Department of Clinical and Surgical Andrology of the Centre of Reproductive Medicine and Andrology, University Hospital of Münster, Germany. Each patient gave written informed consent (ethical approval was obtained from the Ethics Committee of the Medical Faculty of Münster and the State Medical Board no. 2008-090-f-S) and one additional testicular sample for the purpose of this study was obtained. Tissue proportions were snap-frozen or fixed in Bouin’s solution.

### Selection and clinical evaluation of the patient cohort

All patients included in this study underwent full physical examination, hormonal analysis of luteinizing hormone (LH), follicle stimulating hormone (FSH), and testosterone (T), semen analysis (World Health Organization, 2010) and genetic analyses of the karyotype and screening for azoospermia factor (AZF) deletions. All patients had normal karyotypes (46,XY) and no AZF deletions. Exclusion criteria were biopsies of testis with germ cell neoplasia, a history of cryptorchidism of the biopsied testis, and acute infections. To study qualitatively and quantitatively normal spermatogenesis, we collected biopsies from control patients (CTR) that were diagnosed with obstructive azoospermia due to anejaculation (CTR-1) or a congenital bilateral absence of the *vas deferens* due to *CFTR* gene mutations (CTR-2, CTR-3). No sperm was found in the ejaculate but after mechanical dissection of the testicular tissues. To study severely disturbed spermatogenesis, we collected biopsies from cryptozoospermic patients (CZ-1, CZ-2, CZ-3) that were diagnosed with hypergonadotropic oligoasthenoteratozoospermia and presented <0.1 Million sperm in the ejaculate after centrifugation. All CZ patients had elevated FSH levels (Supplementary Table S9).

### Histological analysis of the human testicular biopsies

As a routine diagnostic procedure in our clinic, two testicular biopsies per testis were fixed in Bouin‘s solution, the tissues were washed in 70% ethanol, embedded in paraffin, and sectioned at 5 µm. For histological evaluation, two independent testicular sections from each testis were stained with periodic acid-Schiff (PAS)/ hematoxylin and were evaluated based on the Bergmann and Kliesch scoring method (Bergmann & Kliesch, 2010) as previously described (Siebert-Kuss *et al*, 2023).

### Preparation of single-cell suspensions from testicular biopsies

For the extraction of pure germ cell subtypes, testicular biopsies were digested into a single-cell suspension as previously published (Di Persio et al, 2021). The digestion was based on mechanically chopping up the testicular tissue with a sterile blade into ∼ 1 mm^3^ pieces and a two-step enzymatic incubation, first, with MEMα (ThermoFisher scientific, Gibco, Cat# 22561021) with 1 mg/ml collagenase IA (Merck/Sigma Aldrich, Cat# C9891) at 37 °C for 10 min and, second, with Hank’s balanced salt solution (HBSS) containing 4 mg/ml trypsin (ThermoFisher scientific, Gibco, Cat# 27250018) and 2.2 mg/ml of DNase I (Merck/Sigma Aldrich, Cat# DN25) at 37 °C for 8 – 10 min and strong pipetting in between. Each enzymatic reaction was stopped by adding MEMα supplemented with 10% fetal bovine serum (FBS) (Merck, Cat# S0615) and 1% Penicillin/Streptomycin (ThermoFisher scientific, Gibco, Cat# 15140-148) and the supernatant discarded after centrifugation. Finally, cells were washed three times and the cell suspension then eliminated from erythrocytes by incubation in haemolysis buffer (0.83% NH4Cl solution) for three minutes. The reaction was stopped as described above. Cell debris was removed by filtering the cell suspension through a 70 µm sterile CellTrics® filter (Sysmex). We used the trypan blue exclusion method for quantification of the obtained cell numbers. We incubated the cells (1 million cells / 1 ml HBSS) obtained after digestions with 1 µl Near-IR fluorescent reactive dyeLIVE/DEAD Fixable Dead Cell Stain Kit (Invitrogen; Cat: L34961; Ex: 633/635nm; Em: 775nm) for 30 min on ice and stopped the reaction by addition of 1 ml HBSS supplemented with 5% FBS. After centrifugation, unspecific antibody binding sites were blocked by incubation of the cells with HBSS containing 5% FBS for 45 min on ice. Following centrifugation, cells were fixed in Fix and Perm solution A for 30 min at room temperature and the reaction stopped as outlined above. After centrifugation, cell membranes were permeabilized by incubation with Fix and Perm solution B for 30 min at room temperature. After centrifugation 20% of cells were incubated with fluorophores-conjugated, unspecific immunoglobulin G (IgG) as negative controls, namely mIgG-AI647 (1:200, BioLegend, Cat# 400130), mIgG-Dy550 (1:20, Biolegend Cat#400166) and mIgG-Dy488 (1:20, BioLegend Cat#400166). The remaining cells were incubated with fluorophores-conjugated primary antibodies at room temperature for 1 h against DMRT1-AI647 (1:200, Santa Cruz, Cat# sc-377167 AF647), MAGEA4-Dy550 (1:20, Prof. G. C. Spagnoli, University Hospital of Basel, CH, conjugated) and UTF1-Dy488 (1:20, Merck/Millipore, Cat# MAB4337, conjugated). We used the Dylight^TM^ 488 and 550 labeling kits (ThermoFisher, Cat# 53025 and CAT#84531) for conjugation of the IgGs, MAGEA4 and UTF1 antibodies, respectively. The antibody binding reaction was stopped by adding HBSS supplemented with 5% FBS. After centrifugation, cells were resuspended in HBSS containing 5% FBS and Hoechst (NucBlue® Live ReadyProbes® Reagent Protocol, R37605, Thermo Fisher) was added to distinguish the DNA contents of the cells.

### Fluorescence activated cell sorting analyses for isolation of human male germ cells

For extraction of the different germ cell fractions, we applied a multi-parameter fluorescence activated cell sorting (FACS) strategy on the BD FACS Aria Fusion with FACSDiva software (v 8.02). We gated for cells and live cells based on the Near-IR fluorescent reactive dye LIVE/DEAD^TM^ (LIVE/DEAD™ Fixable Near-IR Dead Cell Stain Kit, Invitrogen, L10119). Spermatogonia were identified and gated within the diploid (2C) cells positive for the pan-spermatogonial marker MAGEA4. The 2C/ MAGEA4+ cells were further divided into undifferentiated spermatogonia (Undiff) and differentiating spermatogonia (Diff) by gating for UTF1^+^/ DMRT1^−^ and UTF1^−^/ DMRT1^+^ cells, respectively. Primary spermatocytes and spermatids were isolated and gated based on their DNA content for 4C and 1C cells, respectively. Cells were sorted with a 85 µm nozzle and collected in 200 µl HBSS containing 5% FBS. FACS data were analyzed using the FlowJo software (v 10.8.1).

### DNA isolation and enzymatic conversion of sorted testicular germ cells

DNA was isolated from fixed and sorted cells using the MasterPure DNA purification kit (MC85200, Lucigen, LGC Ltd, Teddington, UK) following the manufacturer’s protocol. We incubated the cells for 20 min at 90 °C prior to incubation with proteinase K at 65 °C for 1 h. DNA concentration was measured using Qubit® dsDNA HS Assay Kit (life technologies) and a fluorescence plate reader (FLUOstar Omega, BMG Labtech, Germany). Enzymatic conversion was performed using the NEBNext Enzymatic Methyl-seq Conversion Module (New England BioLabs, Cat#E7125S) according to the manufacturer’s protocol.

### Whole-genome enzymatic methylation sequencing

EM-seq libraries were prepared from sorted testicular germ cells of CTR and CZ samples (n=21) using 10-200 ng of DNA supplemented with 0.01% Lambda-/ 0.001% PUC-DNA. DNA was fragmented using the Bioruptor with 3 cycles of 30s on/ 90s off, and the library was prepared with 4-8 PCR cycles. The libraries were sequenced in a NovaSeq6000 instrument using a paired-end 2×150bp protocol and aiming for 80 Gb/sample.

### Data processing and EM-seq data analysis

We processed the raw sequencing data using the wg-blimp pipeline (v 0.10.0) (Wöste *et al*, 2020) (Supplementary Table S10). Originally designed as a pipeline for analyzing whole-genome-bisulfite-sequencing data, wg-blimp is also capable to handle enzymatic sequencing data, which shares the same raw data format but offers improved sequencing accuracy and reliability (Feng *et al*, 2020). wg-blimp incorporates the well-established algorithms for tasks such as bwa-meth for alignment (Li, 2013), MultiQC for quality control (Ewels *et al*, 2016), MethylDackl (v 0.6.1) (Ryan, 2021) for methylation calling, and camel (v 0.4.7) (Schröder, 2018) and metilene (v 0.2-8) (Jühling *et al*, 2016) for identifying DMRs. All data were aligned against the GRCh38.p7 reference genome.

To visualize the distribution of DNA methylation across various functional regions, we utilized the annotation sources provided by wg-blimp for hg38 alignment, which includes gtf-annotation and masked repeats. These alignments were imported into R 4.2.1 (R Core Team, 2022) using the *“rtracklayer”* (v 1.58.0) (Lawrence *et al*, 2009), while the identification of methylated regions of interest was performed using the *“GenomicRanges”* package (v 1.50.2) (Lawrence *et al*, 2013). Statistical analyses and graph plotting were performed using R packages, namely *“stats”* (v 4.2.1), *“ggplot2”* (v 3.3.6) (Wickham, 2016), and *“introdataviz*” (v 0.0.0.9003) (Nordmann *et al*, 2022). For the analysis of methylation distribution, we considered only CpG sites with a minimum coverage of 5.

Principal component analysis (PCA) was conducted on the methylation values, which were obtained from the wg-blimp software using the tool MethylDackl. The PCA analysis was conducted on 2521 CpG sites within the 50 known imprinted control regions (ICRs) (Monk *et al*, 2018), where all samples exhibited at least 5 x coverage.

### DMR analyses

DMR calling was conducted using the wg-blimp software. The following criteria for DMR identification were applied: A minimum coverage of 5 per CpG loci, covering at least 5 CpG sites within a DMR, showing at least 20% methylation difference in the compared groups, and a maximum mean difference of ≤ 30% within each group for the CTR samples and ≤15% for the CZ samples, to reduce the influence of genetic variability between patients. For metilene, a threshold of qD<D0.05 was set, whereas camel uses t-statistics for verification.

Positive or negative enrichment of DMRs within specific genomic regions was assessed using permutation tests. The regions of interest were annotated using R 4.3.0 (R Core Team, 2023) with the packages “stats” (v 4.3.0), "*annotatR*" (Cavalcante & Sartor, 2017) and *"TxDb.Hsapiens.UCSC.hg38.knownGene"* (v 3.17.0) (Team BC & Maintainer BP, 2019), while the masked repeats were annotated using the annotation provided by wg-blimp. Permutation tests were performed using *"regioneR"* (Gel *et al*, 2016) and "*GenomicRanges*" using a Bonferroni correction (α=0.00019). Differences were quantified using z-scores and p-values provided by the permutation test with 10,000 iterations in *"regioneR"*.

To evaluate the average expression and percentage of cells expressing the DMR associated genes identified in this study, we used a previously published dataset (Di Persio *et al*, 2021b). Evaluation was performed using Seurat (Stuart *et al*, 2019; Hao *et al*, 2021). For better comparison to our dataset, we summarized the spermatocyte and spermatid cells from the scRNA-seq dataset together. Germ cell type-specific genes for undifferentiated spermatogonia, differentiating spermatogonia, spermatocytes, and spermatids were extracted based on differential gene expression analysis using MAST (Finak *et al*, 2015). Filtration criteria for germ cell type-specific genes were: log fold change threshold of ≥ 0.5, FDR-corrected p-value below 0.01 and expression in at least 25% of the cells of one comparison group.

### GO term analyses

*“ChIPseeker”* (Yu *et al*, 2015) was used to retrieve a comprehensive gene list of overlapping genes, gene promoters (TSS ± 1000 bp) and flanking genes (putative regulatory sites, 5000 bp) from our DMRs. These comprehensive DMR associated gene lists were then analyzed for GO term enrichment for BP, MF, and CC using *“clusterProfiler”* (Wu *et al*, 2021) and the enrichR database (Xie *et al*, 2021). P-Value was adjusted for multiple testing with Benjamini-Hochberg correction.

### Motif analysis

To identify enriched known motifs of genes and TFs within the DMRs, we used HOMER (v 4.11) (Heinz *et al*, 2010). This tool was utilized with the default parameter of the fragments size of 200 bp and with the “-mask” parameter to use the repeat-masked sequences. Notably, HOMER uses regions with the same GC-content distribution as control.

### Retrieval of public datasets

We downloaded the GSE40195 dataset from the GEO database, which contains ChIP-seq data of enriched regions for retained histones (H3.3, H3K14ac, H3K27ac, H3K36me3, H3K4me1 and H3K9me3) in human sperm. The data was converted to GRCh38/hg19 using “*rtracklayer”*.

### Statistical analyses

Statistical analyses was performed as described in sections for data processing and EM-seq data analyses, DMR analysis, genomic annotation of the DMRs, and motif analysis.

## Data availability

EM-seq data has been deposited in the European Genome-Phenome Archive under EGAS00001007449.

