## Supplementary figures and images for "Genome-wide DNA methylation changes in human spermatogenesis"

### Supplementary Fig. S1

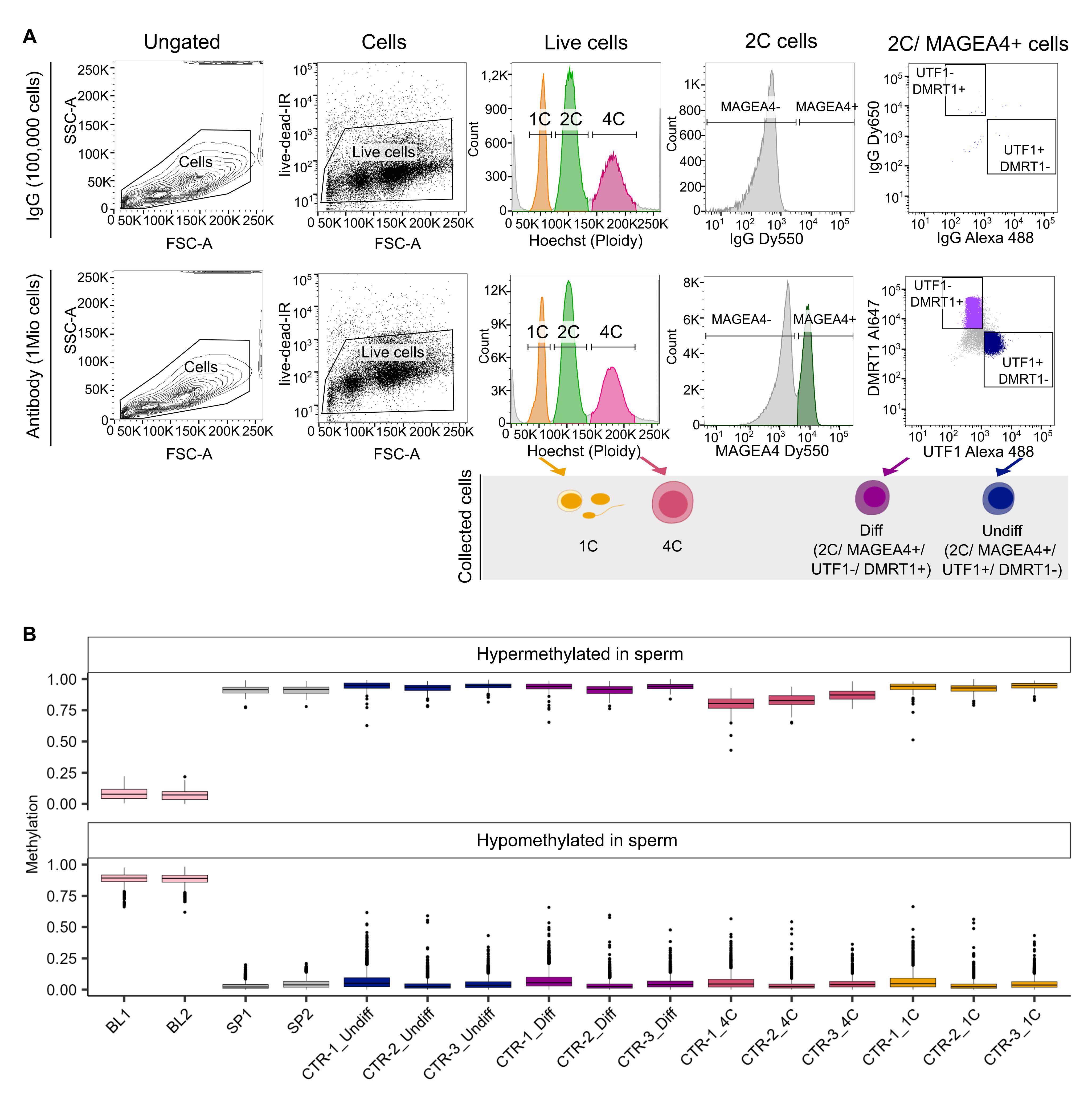

### Supplementary Fig. S2

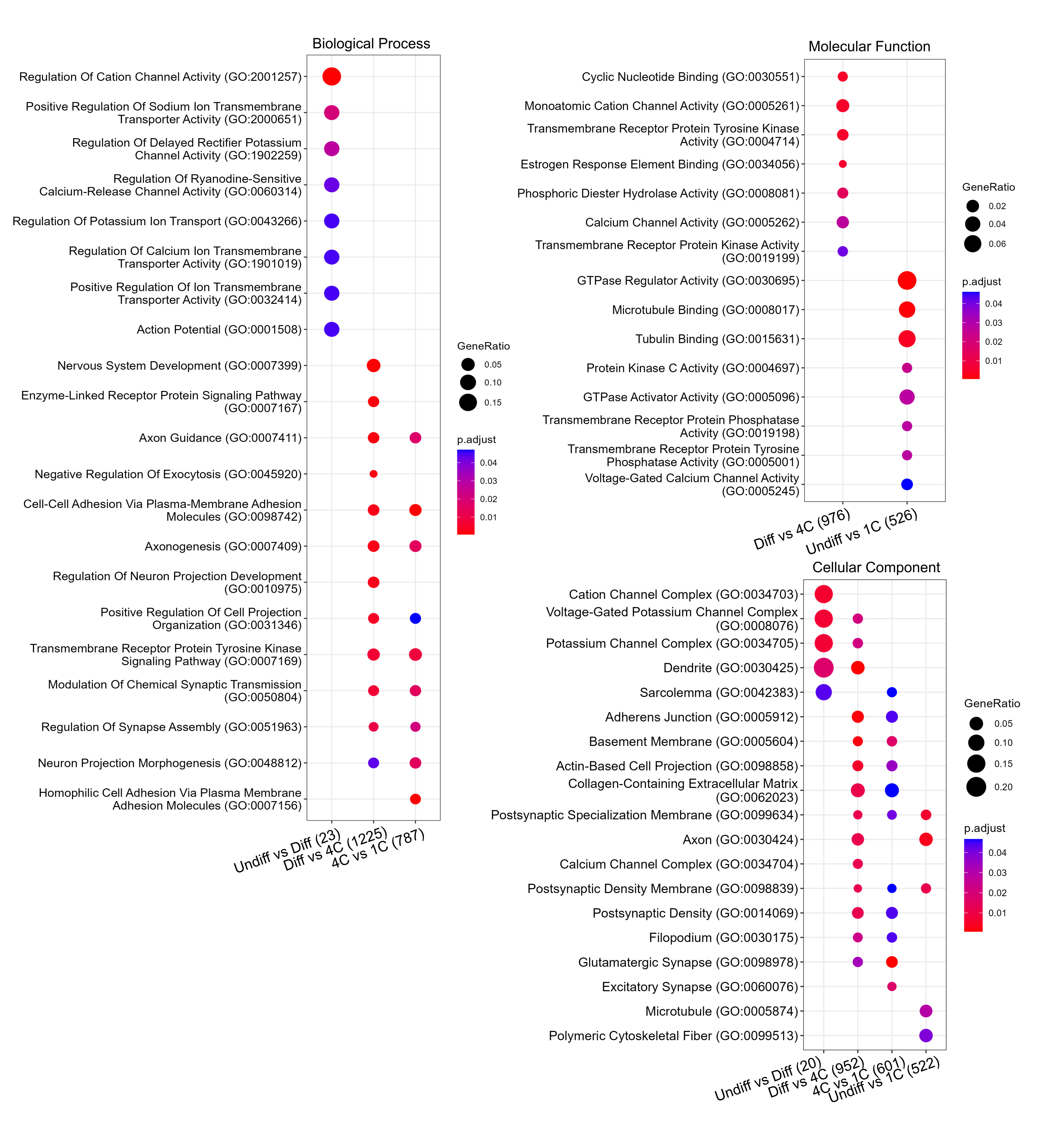

### Supplementary Fig. S3

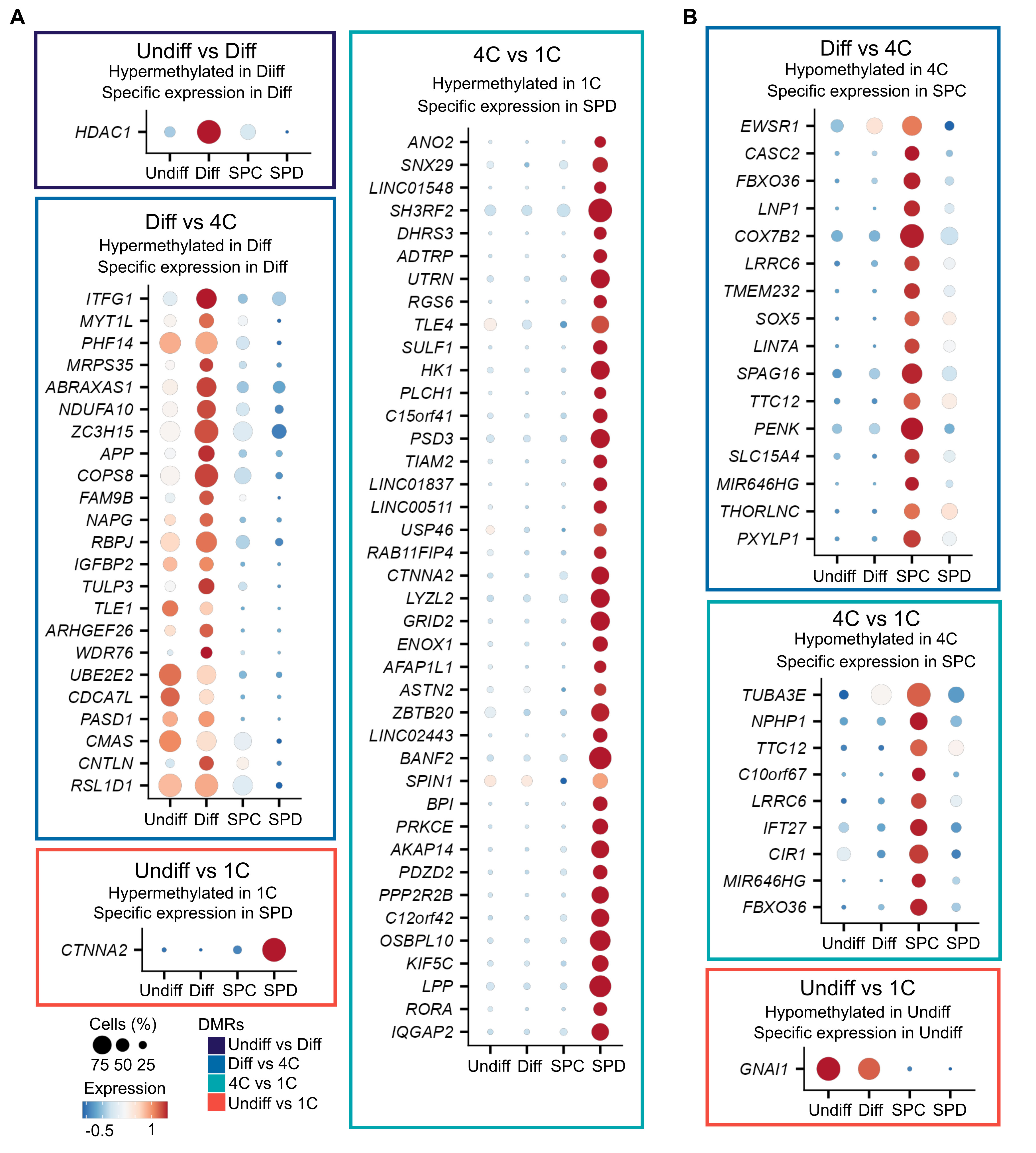

### Supplementary Fig. S4

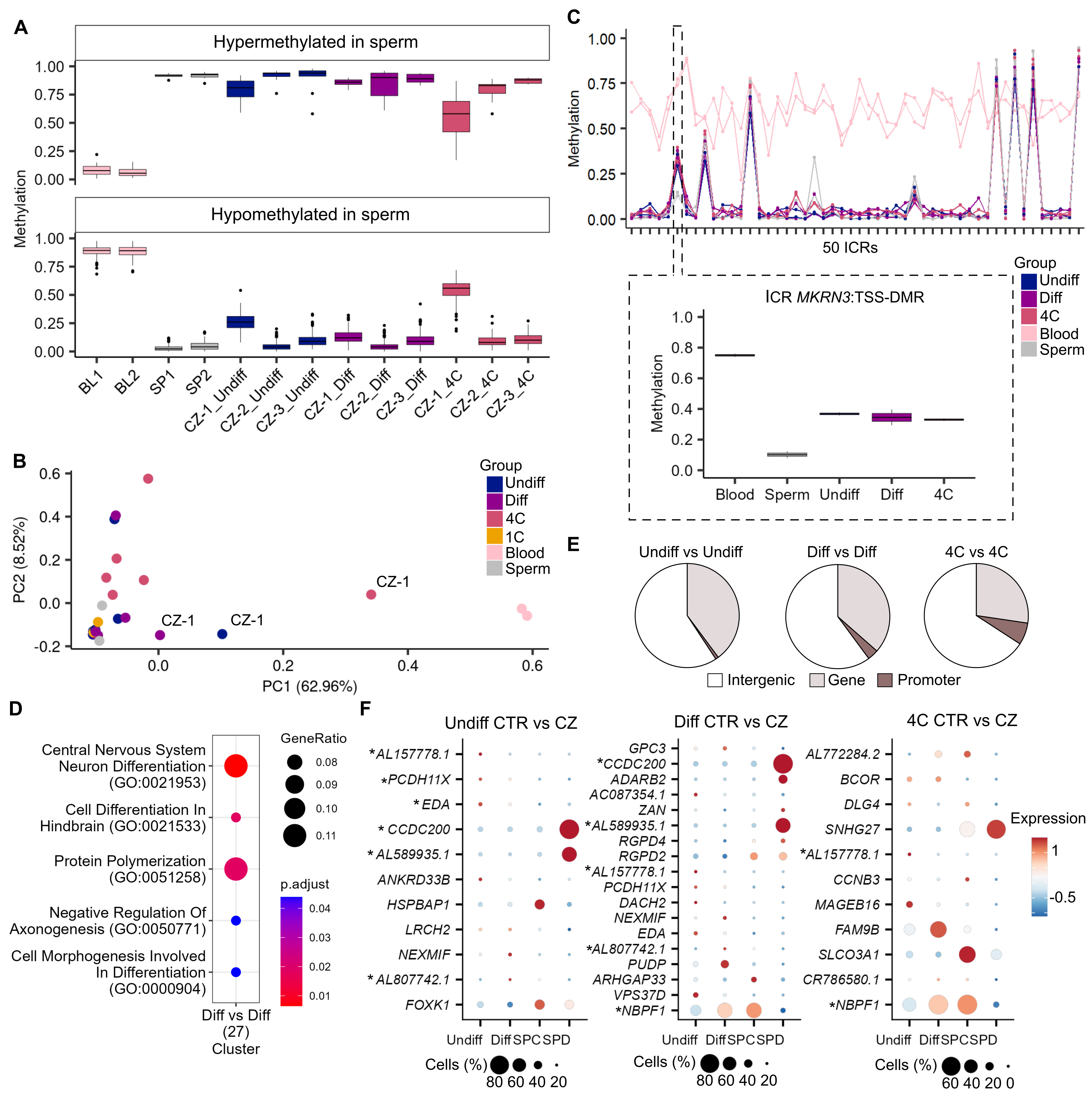

### Supplementary Fig. S5

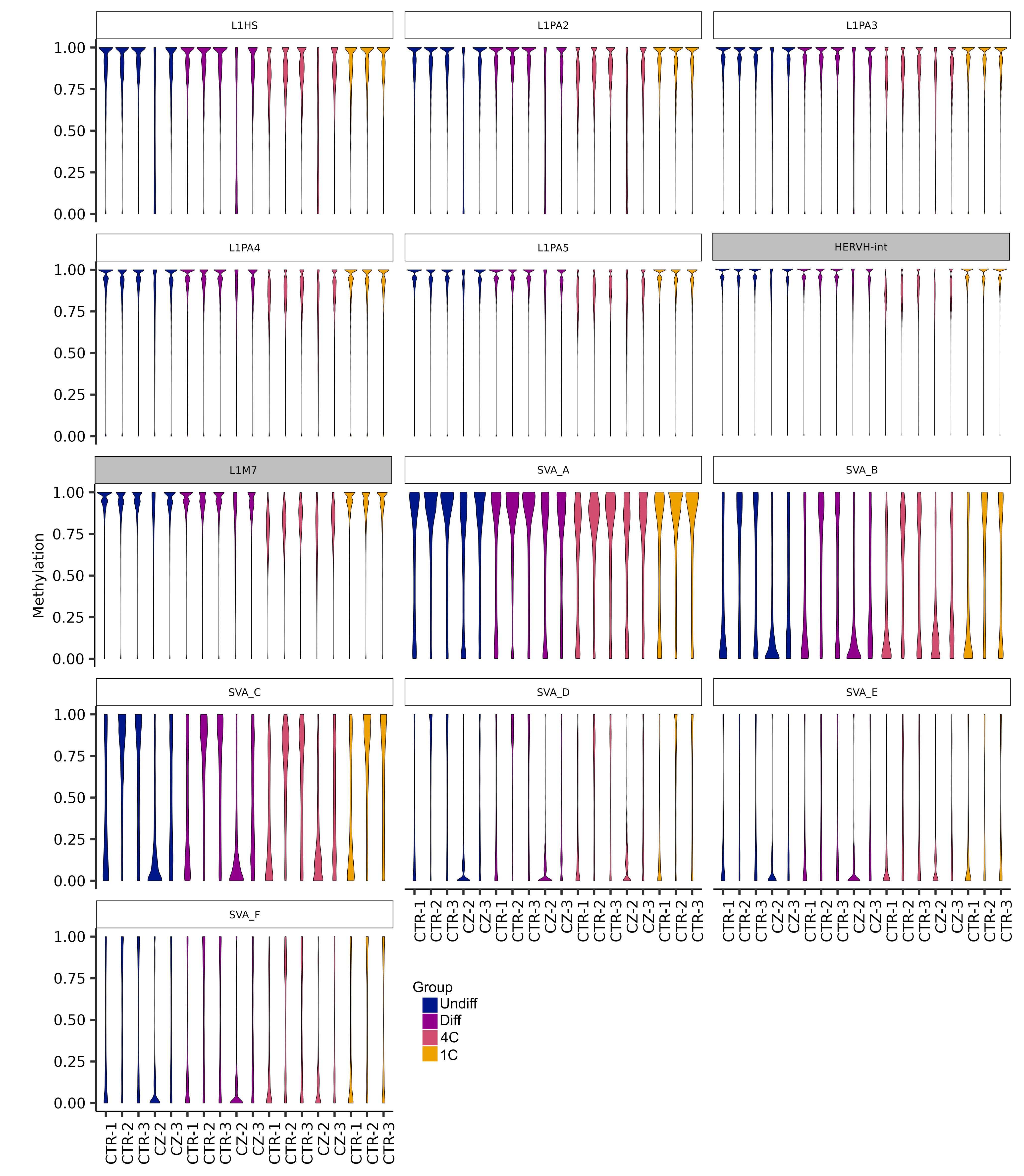
